# Genome sequencing confirms cryptic diversity and potentially adaptive changes in gene content of Lepidoptera-infecting Apicomplexa

**DOI:** 10.1101/2025.05.29.656805

**Authors:** Andrew J. Mongue, Peter Kuperus, Astrid T. Groot

**Affiliations:** Department of Entomology and Nematology, University of Florida, USA; Institute for Biodiversity and Ecosystem Dynamics, University of Amsterdam, Netherlands

## Abstract

Apicomplexan parasites of insects represent a largely unexplored branch of the tree of life. Their small size and lack of informative phenotypic traits can obscure their diversity and many lineage-specific adaptations to different host species. To better understand how these parasites have evolved, we have sequenced and annotated the genome of a neogregarine parasite of moths including *Helicoverpa armigera*. This parasite, which we term *Helicoverpa-*infecting *Ophryocystis*, has a tiny 7 Megabase genome with only 2,200 protein-coding genes. Using phylogenomic methods, we place it as the closest currently sequenced relative to *Ophryocystis elektroscirrha* and its sister lineage, both of which infect milkweed butterflies. We then explore evolution of the ATPase gene family, which is known to be reduced in other *Ophryocystis* lineages. *Helicoverpa-*infecting *Ophryocystis* has more ATPase genes than other *Ophryocystis*, which may relate to the interactions of parasites with phytochemistry encountered in their insect hosts. These results demonstrate both the amount of uncertainty remaining in the insect-infecting Apicomplexa as well as the utility of applying modern genomic approaches to their study.

## Introduction

Apicomplexa are ancient and diverse unicellular eukaryotic parasites (Levine, 2018). Members of the phylum infect most multicellular animals, including both vertebrates like humans, (Abrahamsen et al., 2004; Gardner et al., 2002; Levine, 2018) and invertebrates (Gao et al., 2020; McLaughlin & Myers, 1970; Templeton et al., 2010). This evolutionary longevity and host breadth makes them intriguing targets for evolutionary genetic study: what molecular and genomic features are required to exploit so many hosts across the tree of life?

Some species, such as the causal parasite of malaria, *Plasmodium*, have understandably received intensive study (Bushman et al., 2016; Gardner et al., 2002; Krishna et al., 2001). However, much of the diversity remains largely unexplored, especially that of invertebrate-infecting Apicomplexa (Boisard & Florent, 2020). Better understanding of these parasites is important both to make sense of the evolution and diversity of life at a basic level (Boisard & Florent, 2020; DeBarry & Kissinger, 2011) and to develop potential biological controls of pest arthropods (e.g., as control agents of mosquitos, Tseng, 2007). The first step towards these two broader goals is to gain a better molecular understanding of invertebrate-infecting Apicomplexa that are already being used as model systems in the laboratory.

Recent sequencing of two *Ophryocystis* lineages (Mongue et al., 2023) that parasitize milkweed butterflies (Nymphalidae Danainae) has already helped to shed light on the host-parasite interactions of this system. In particular, larvae of butterflies in the genus *Danaus* are specialists on milkweed plants in the genus *Asclepias*, which contain toxic cardiac glycoside phytochemicals (Agrawal et al., 2012; Brower & Fink, 1985). These compounds block the function of sodium/potassium pump ATPases in most organisms (Matsui & Schwartz, 1968), but monarchs possess amino acid adaptations that make them resistant to cardiac glycosides (Aardema & Andolfatto, 2016). Given the toxic effects of milkweed phytochemicals on ATPase function generally (Kelly et al., 1985; Matsui & Schwartz, 1968), and the demonstrated inhibitory effect of milkweed chemistry on apicomplexan parasite growth (de Roode et al., 2008; De Roode et al., 2011), an intuitive hypothesis was that the milkweed phytochemicals would inhibit ATPase. However, in sequencing the genome of *Ophryocystis*, the authors found that these parasites lack sodium pump ATPases entirely (Mongue et al., 2023). This unexpected finding led to the hypothesis that exposure to milkweed phytochemicals could lead to the loss of ATPases, although it was difficult to draw conclusions about parasite adaptation with only a pair of sequences from milkweed butterfly parasites and no relatives to act as comparisons.

Here we report the genome sequence of another *Ophryocystis* parasite of moths in the genus *Helicoverpa* (Lepidoptera: Noctuidae). This parasite has been described as *Ophryocystis elektroscirrha*-like (Gao et al., 2020) and is visually nearly identical to *O. elektroscirrha* in the most visible oocyst stage. However, unlike *O. elektroscirrha’s* host, *Helicoverpa* moths are generalist crop pests whose diet does not include cardiac glycoside producing plants (Nasreen & Mustafa, 2000; Zalucki et al., 1986). As such, their parasites should not be exposed to the selective pressures of ATPase inhibiting phytochemicals. The *Ophryocystis* parasite of *Helicoverpa* thus makes a useful outgroup to understand the genetics of ATPases and how they compare to *O. elektroscirrha*. To explore the evolutionary genetics of ATPases, as well as to generate resources for another insect-infecting apicomplexan model, we present the genome of a previously unsequenced *Ophryocystis* species and explore the molecular evolution of ATPases.

## Methods

### Sample collection and DNA extraction

Oocyst purification was performed on *Chloridea virescens* moths that had been infected with *Ophryocystis elektroscirrha*-like parasite, collected originally from *Helicoverpa armigera* (see Gao et al. 2020). To remove debris and isolate oocyst, wings were removed from dead, infected moths. Groups of 10 moths were pooled into 50 mL tubes (eight sets, totalling 80 moths). Each tube was supplemented with 40 mL sterile water and 1 mL Dreft detergent solution and subjected to two rounds of 10-minute sonication. Samples were then manually shaken for 10 minutes, minimizing foam formation. Large insect fragments were removed before the tube was centrifuged at 1,000 rpm for 2 minutes in a fixed-angle rotor, and oocysts were collected from the black sediment layer. The oocyst-rich black layer (Figure 1, left) was collected into a 2 mL tube, washed twice with 1 mL water/Dreft solution, and vortexed before centrifuged at 10,000 rpm for 4 minutes. The black oocyst pellet was retained, while the brown layer of moth scales was removed by pipetting. Oocysts were air-dried and resuspended in 80 μL TE buffer. Oocyst integrity and purity were assessed microscopically (∼2 μL per sample, Figure 1, middle). DNA extraction was performed using a modified CTAB protocol (Doyle and Doyle, 1987). First, a total of 10 to 20 zirconium beads (1 mm) were added to the spore suspension, followed by homogenization using a Precellys system (6,800 rpm). Oocyst disruption was confirmed microscopically (Figure 1, right).

**Figure 1.**
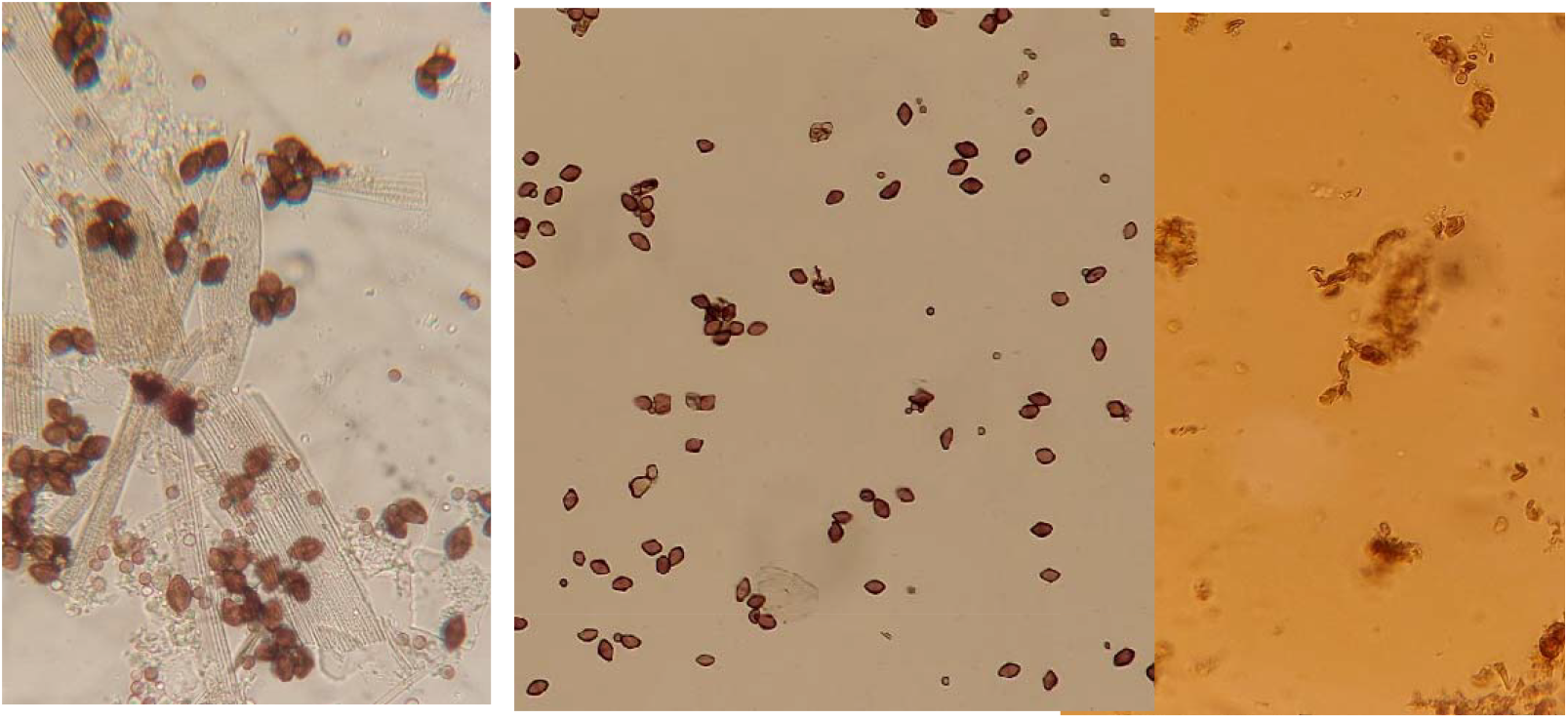
Oocyst purification for DNA extraction. **Left:** Oocyst-rich suspension collected from *C. virescens* adults. **Middle:** Cleaned oocyst suspension. **Right:** Disrupted oocysts for DNA extraction.

For DNA extraction, 600 μL of 2% CTAB solution and 6 μL Proteinase K (Thermo Scientific, 20 mg/mL) were added, followed by incubation at 56°C for 1 hour. DNA was purified by adding chloroform:isoamyl alcohol (24:1), vortexing, and centrifuging at 14,000 rpm for 8 minutes. The upper aqueous phase transferred to a new tube, and the extraction was repeated. DNA was precipitated with 96% ethanol and incubated at - 20°C for 30 minutes, followed by centrifugation at 14,000 rpm for 10 minutes at 10°C. The DNA pellet was washed with 80% ethanol, centrifuged under the same conditions, and resuspended in 50 μL TE buffer containing 2 μL 5M NaCl and 2 μL RNase (1 mg/mL). Samples were incubated at 37°C for 30 minutes, after which DNA was re-precipitated using 150 μL 96% ethanol, incubated at -20°C for 30 minutes, and centrifuged again. The pellet was washed with 200 μL 80% ethanol, air-dried, and resuspended in 25 μL sterile water. DNA concentration and purity were assessed using a Implen nanophotometer (N60 Touch), yielding 8.9 ng/μL (260/280 = 1.9; 260/230 = 1.6).

### Sequencing and genome assembly

Library preparation started with 204 ng of DNA and followed the Oxford Nanopore genomic-dna-by-ligation-sqk-lsk110 protocol. After the final adapter ligation was performed, DNA concentration was measured on a nanophotometer (2.6 ng/μL; 260/280 = 1.3; 260/230 = 0.4). Sequencing was performed on an Oxford Nanopore Flongle flow cell (FLO-FLG001) loaded with 13 ng library prepped DNA and run for 24 hours.

A minimum quality filter was implemented to remove low quality (<Q10) reads from the generated data. Genome assembly was performed using Geneious Prime (version 2023.0.4) (https://www.geneious.com/prime/) with the Flye and Minimap2 plugins. The flye assembler was run in –nano-raw mode with an estimated genome size parameter of 10m, based on results from existing gregarine genomes.

### Phylogenomics

To assess evolutionary relationships, a phylogenomics pipeline that was previously successfully implemented in insects by Mongue et al. (2024) was employed for Apicomplexa. In brief, the genome assemblies listed in **Table 1** were obtained from online databases and BUSCO v4.14 (Manni et al., 2021) was run in genome mode using the apicomplexa_odb10 dataset to identify single copy orthologs in each species. Next, scripts from the BUSCO_phylogenomics git repo (https://github.com/jamiemcg/BUSCO_phylogenomics) were used to parse complete single copy orthologs found in at least 70% of the sampled species. Subsequently, FastTree v2.1.11 (Price et al., 2009) was used to construct gene trees for each orthologous protein with the JTT model of molecular evolution (Jones et al., 1992) and the CAT approximation for variable rates at each site (Stamatakis, 2006). Finally, ASTRAL (Zhang et al., 2018) was used to generate a single consensus species tree with local quartet posterior probabilities for support (Sayyari & Mirarab, 2016).The tree was manually rooted with a dinoflagellate, *Symbiodinium microadriaticum*, as the outgroup.

**Table 1.** Genomic resources for phylogenomic analyses. For each species we list the data accession and publication, where available.

| Species | Accession | Reference |
| --- | --- | --- |
| *Ascogregarina taiwanensis* | PRJNA27765 | (Templeton et al., 2010) |
| *Cryptosporidium meleagridis* | PRJNA1022047 | (Penumarthi et al., 2024) |
| *Cryptosporidium parvum* | PRJNA144 | (Abrahamsen et al., 2004) |
| *Eimeria praecox* | PRJEB71489 | (Blake et al., 2024) |
| *Gregarina niphandrodes* | PRJNA259233 | Unpublished |
| *Ophryocystis elektroscirrha* | PRJNA906508 | (Mongue et al., 2023) |
| *Ophryocystis elektroscirrha*-like | https://doi.org/10.5281/zenodo.7817702 | (Mongue et al., 2023) |
| *Helicoverpa*-infecting *Ophryocystis* | [TBD] | This publication |
| *Plasmodium falciparum* | PRJNA13173 | (Gardner et al., 2002) |
| *Plasmodium yoelii* | PRJEB5706 | (Otto et al., 2014) |
| *Porospora gigantea* A | PRJNA734792 | (Boisard & Florent, 2020) |
| *Porospora gigantea* B | PRJNA734792 | (Boisard & Florent, 2020) |
| *Sarcocystis neurona* | PRJNA227351 | (Dangoudoubiyam et al., 2024) |
| *Symbiodinium microadriaticum* | PRJNA292355 | (Aranda et al., 2016) |
| *Theileria orientalis* | PRJDA49463 | (Hayashida et al., 2012) |

### Gene annotation

A gene annotation method was chosen that has proven successful for other *Ophryocystis* genomes (Mongue et al., 2023). Specifically, the GenSAS pipeline (Humann et al., 2019) was used to generate gene models as follows. To start, repetitive sequences were identified with RepeatMasker v4.1.1 (Smit et al., 2019) and masked with RepeatModeler v2.0.1 (Flynn et al., 2020) and in GenSAS (Humann et al., 2019). Based on previous success, GeneMarkES v4.48 (Ter-Hovhannisyan et al., 2008) was used to annotate genes in the repeat-masked assembly, then BUSCO v5.2.2 (Manni et al., 2021) was used to query the annotation with the apicomplexa_odb10 dataset of putatively conserved orthologs.

### General orthology analysis

With the sequencing and annotation of this parasite, there are now three genomes and annotations from lineages in Ophyrocystidae, enough information to begin exploring overall patterns of gene conservation in this family. To do so, ProteinOrtho v5.11 (Lechner et al., 2011) was used to search for orthologs between *Helicoverpa*-infecting *Ophryocystis* and the two existing *Ophryocystis* lineages from milkweed butterflies. Basic summary statistics of gene conservation are presented in the results.

### ATPase gene family analysis

Because the *Helicoverpa* parasite represents a closer relative to other *Ophryocystis* lineages than other sequenced Apicomplexa and its host does not associate with cardiac glycosides, the ATPase genes from this genome serve as interesting negative control contrasts to those described from *O. elektroscirrha* and its sister lineage (Mongue et al., 2023). To investigate the ATPase genes, we followed our previous procedure (Mongue et al. (2023) by querying the new gene annotation with the amino acid sequence of a known ATPase gene (GenBank accession: CCJ05450.1) using BLAST+ v2.9.0 (Camacho et al., 2009). ATPase function was confirmed via reciprocal BLAST on NCBI. Once confirmed, these new sequences were added to a database of primarily apicomplexan ATPases that was initially compiled by Lehane et al. (2019) and further developed by Mongue et al. (2023). These sequences were aligned with CLUSTAL Omega’s web-hosted tool (Sievers & Higgins, 2014) and manually trimmed the output to ∼260 conserved amino acids. To generate a phylogenetic tree, this trimmed alignment was passed to the *phangorn* package in R (Schliep, 2011) using the LG+G(4)+I model of substitution and 100 bootstrap replicates to give confidence. With this final phylogenetic tree, the function of newly identified ATPases was inferred based on sequence similarity to those with existing functional annotations.

## Results

### Genome assembly

We generated 249,580 Oxford Nanopore reads with an N50 of ∼2,100 Kb, totaling 304.9 Mb of sequence data. Of these, 172,880 passed base-calling quality filter (>Q10) and were used for downstream assembly, totaling 172.9 Mb. Primary assembly stats are found in **Table 2**. Of note, only two contigs had lengths <1,000 bp.

**Table 2.**
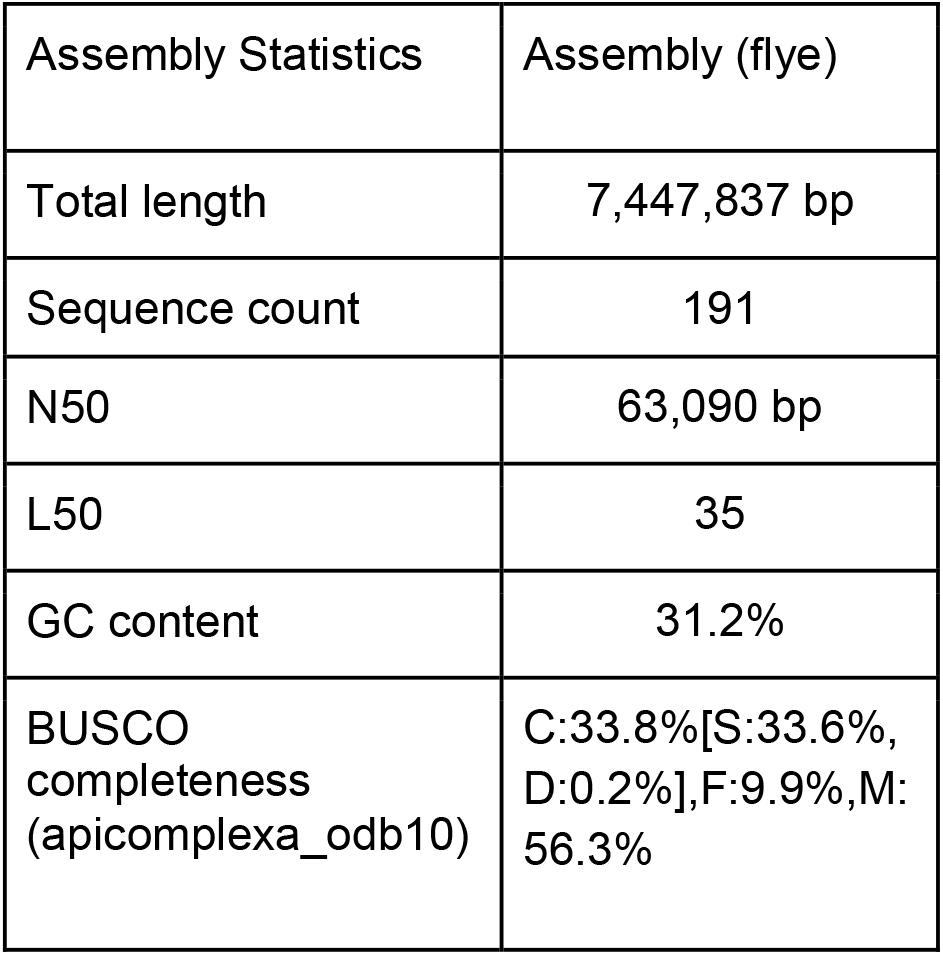
Genome assembly statistics.

### Phylogenomics

As expected, but now confirmed via genomic analysis, the *Ophryocystis* parasite of *H. armigera* is genetically distinct from *O. elektroscirrha* that infects *D. plexippus*, and in fact represents the outgroup to the two lineages of milkweed butterfly infecting *Ophryocystis* (**Figure 1**).

**Figure 1.**
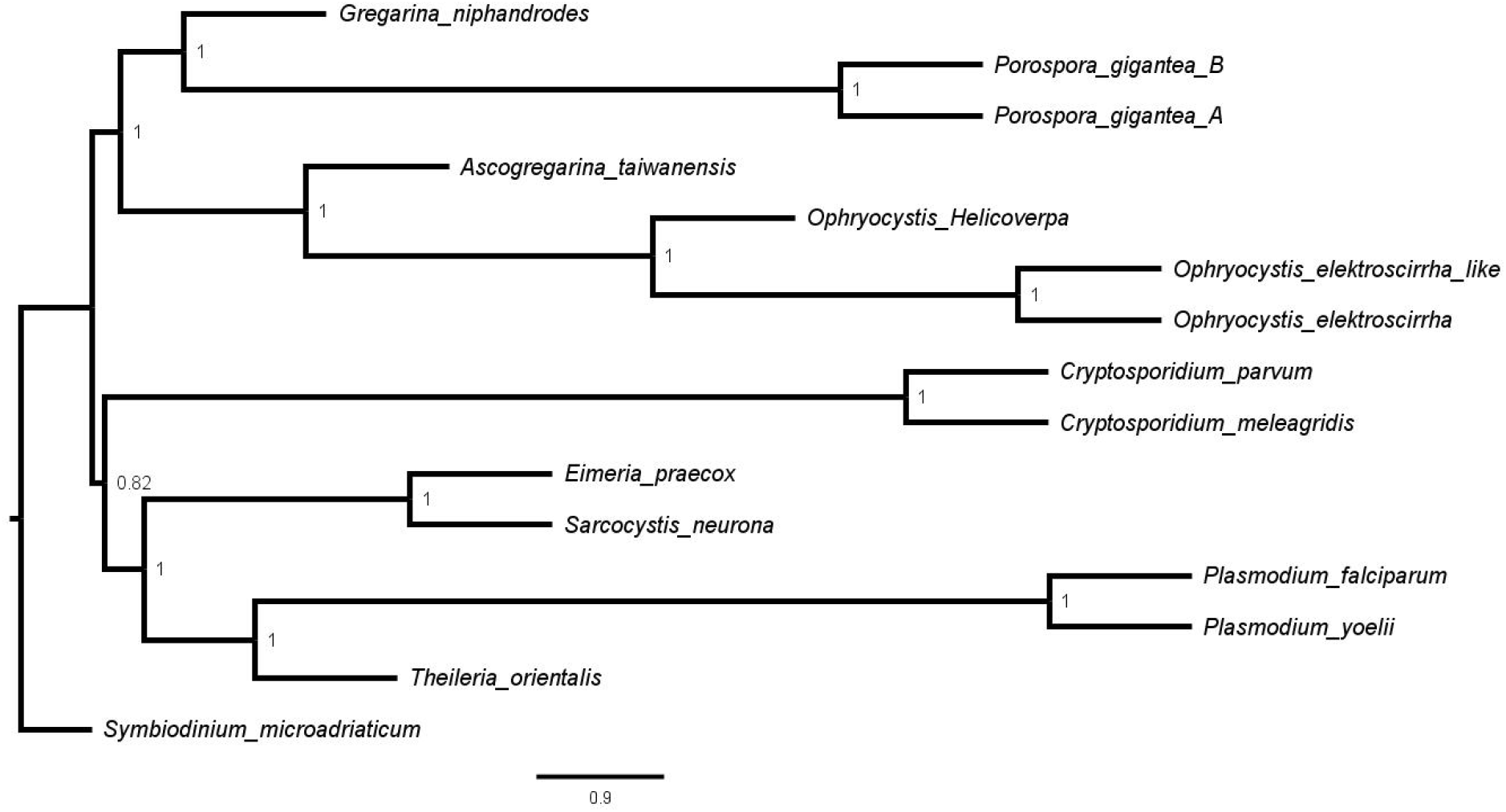
Phylogenetic placement of *Helicoverpa* infecting *Ophryocystis*. *Ophryocystis* parasites of *H. armigera* are sister to the lineages of *Ophryocystis* found in milkweed butterflies. Node labels show posterior quartet support values.

*Ascogregrina taiwnensis*, a parasite of mosquitos (Templeton et al., 2010), is the closest sequenced relative of *Ophryocystis* as currently defined. More generally, all of these genomes fall into a monophyletic clade with other species in the subclass Gregarinasina, all of which are parasites of arthropods.

### Repeat and gene content

As expected of a very small genome, we recovered a very low repeat content (9.74%), which lower even that the ∼12% seen in *O. elektroscirrha* (Mongue et al., 2023). All of these were simple repeat motifs; RepeatModeler found no complex families (e.g. retrotransposons). Likewise, we recovered a mere 2,225 genes encoding 2,177 proteins. This gene set contained 61.2% (n=273) of expected complete and single copy orthologs, 0.7% (n=3) duplicated complete BUSCOs, 3.6% (16) fragmented and 34.5% (n=154) missing BUSCOs. Despite the low absolute numbers, we note that this annotation marks an improvement over the BUSCOs identified directly from the genome, strongly suggesting this annotation is not failing to capture genes in the underlying genome.

### General patterns of orthology

We recovered 2,094 genes conserved in at least two of the three species in some capacity, *i*.*e*. either 1-to-1 or 1-to-many. Starting with pairwise orthology, we found 556 genes conserved in *O. elektroscirrha* and *O. elektroscirrha*-like but absent in *Helicoverpa*-infecting *Ophryocystis*; all of these were 1-to-1 strictly conserved orthologs. Another 489 were found in *Helicoverpa*- infecting *Ophryocystis* and *O. elektroscirrha* but not *O. elektroscirrha*-like, and finally a mere 56 proteins were conserved in *Helicoverpa*-infecting *Ophryocystis* and *O. elektroscirrha*-like but not *O. elektroscirrha*. After accounting for these pairwise orthologs, we identified 993 proteins conserved across all three lineages. Of these, only 15 were 1-to-many orthologs, and in each of those cases, the relationship was a 1-to-1-to-2 orthology, in other words most parsimoniously single lineage-specific gene duplication events. Specifically, we recovered 5 duplications in *Helicoverpa*-infecting *Ophryocystis*, 9 duplicate orthologs in *O. elektroscirrha*, and 1 single duplicate ortholog in *O. elektroscirrha*-like. Simple subtraction of these counts from the total annotated gene sets leaves 633 lineage specific genes in *Helicoverpa*-infecting *Ophryocystis*, 648 unique to *O. elektroscirrha*, and 674 unique to *O. elektroscirrha*-like. Results are summarized in **Table 3**.

**Table 3.** Summary of gene conservation and novelty across three lineages of Ophryocystidae. Greyed out diagonal cells represent self-vs-self comparisons and list the total gene content. For the Vs. all comparison, 993 is the number of 1-to-1-to-1 orthologs, with the parenthetical giving the number of lineage-specific duplicate genes.

| Lineage | Vs.<br>*Helicoverpa*-<br>infecting<br>*Ophryocystis* | Vs. *O.*<br>*elektroscirrha* | Vs. *O.*<br>*elektroscirrha*-<br>like | Vs. all | Lineage<br>specific |
| --- | --- | --- | --- | --- | --- |
| *Helicoverpa*-<br>infecting<br>*Ophryocystis* | 2,177 | 489 | 56 | 993 (+5) | 634 |
| *O. elektroscirrha* | 489 | 2,695 | 556 | 993 (+9) | 648 |
| *O.*<br>*elektroscirrha*-<br>like | 56 | 556 | 2,280 | 993 (+1) | 674 |

### ATPase gene family evolution

In total, we identified four ATPases from *Helicoverpa* infecting *Ophryocystis* (**Figure 2**, closed red triangles). Three of these are orthologous to ATPases found in the genomes of *O. elektroscirrha* and the related *O. elektroscirrha*-like (**Figure 2**, open black triangles). One of these (HeliOph.00g018340.m01) is inferred to be a sarcoplasmic/endoplasmic reticulum Calcium ATPase (SERCA), though with a notably longer branch and lower bootstrap support compared to the other two *Ophryocystis* SERCAs. The other two are orthologous to a pair of *Ophryocystis* ATPases (HeliOph.00g000870.m01and HeliOph.00g012140.m01) that were previously identified as the Plasma Membrane-bound Calcium (PMCA) ATPases. In our current analyses however, these ATPases are confidently placed apart from other PMCAs and showed reciprocal BLAST hits to phospholipid transporting ATPases. For each of the three ATPases above, the gene phylogeny recapitulates the species phylogeny, with the newly sequenced *Ophryocystis* being the outgroup to the other two *Ophryocystis* sequences. Finally, *Helicoverpa* infecting *Ophryocystis* has a fourth ATPase (HeliOph.00g001320.m01) not found in the other *Ophryocystis*. This ATPase is confidently placed within the Plasma Membrane-bound Calcium (PMCA) ATPases of *Cryptosporidium* and other Apicomplexa (**Figure 2**).

**Figure 2.**
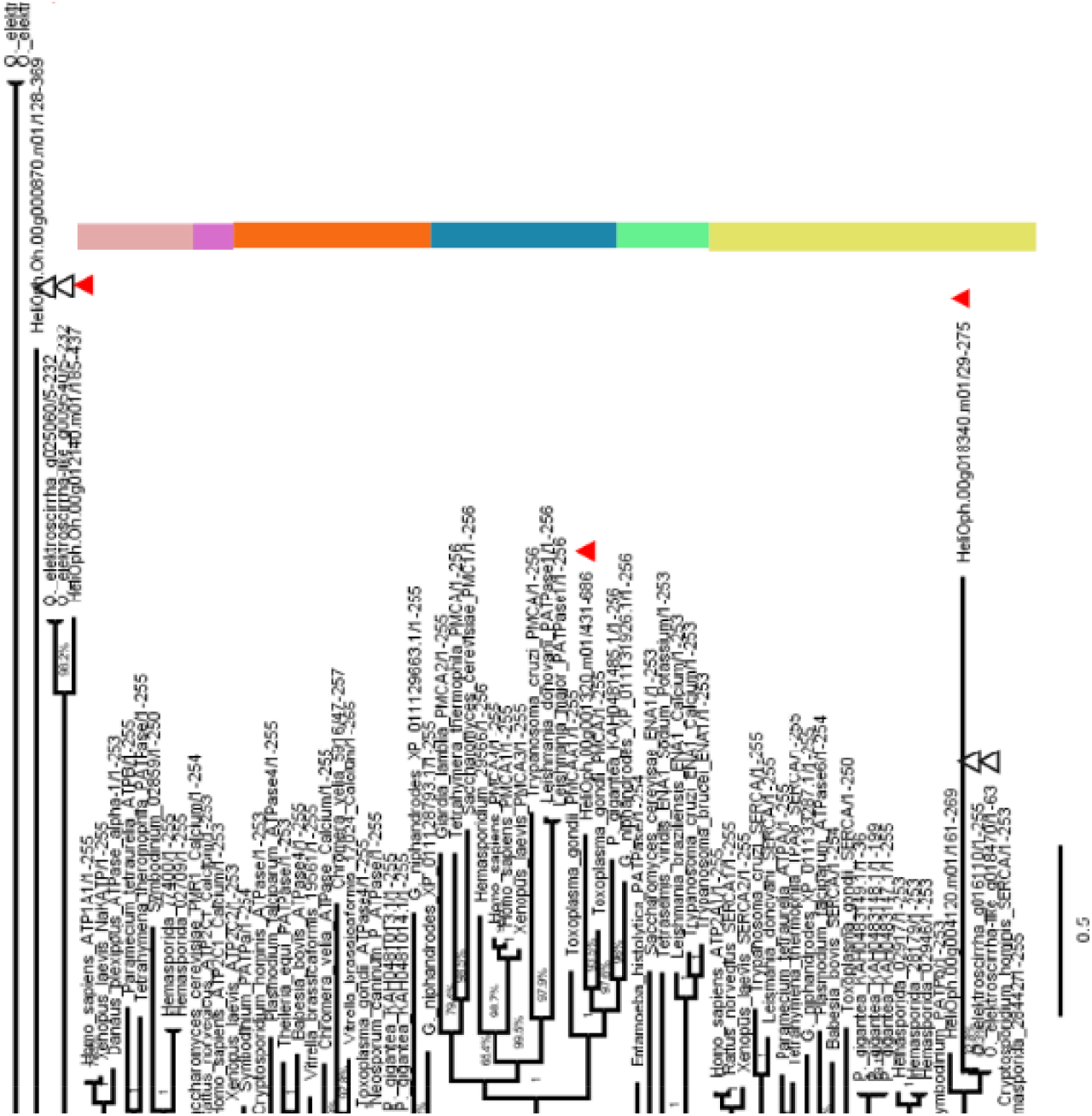
Maximum likelihood tree of ATPase family genes across Apicomplexa. ATPase genes from *Helicoverpa* infecting *Ophryocystis* are denoted with filled red triangles, those of other *Ophryocystis* with open black triangles. Functions are inferred from relation to other apicomplexan ATPases with known function.

## Discussion

We offer the first sequence of *Helicoverpa*-infecting *Ophryocystis*, an apicomplexan parasite of moths in the genus *Helicoverpa*. This parasite has been called “*Ophryocystis elektroscirrha*-like”, although previous phylogenetic analysis of 18S rRNA placed it in a separate clade from other similar parasites of *O. elektroscirrha sensu stricto* (Gao et al., 2020). Our present analyses confirm the distinction between these visually similar parasites at the genomic and genetic level.

### *Comparative genomics of* Ophryocystis

To start with, the genome of *Helicoverpa*-infecting *Ophryocystis* spans only 7.1 Mb, roughly 1.7 Mb smaller than the two other published *Ophryocystis* genomes and with fewer protein coding genes (Mongue et al., 2023). Although we lack independent estimates for genome sizes in this clade (e.g. flow cytometry), the consistency of reports of small genomes in other *Ophryocystis* (Mongue et al., 2023) and their closest sequenced outgroup *Ascogregarina taiwanensis* (a mere 6.1 Mb, Templeton et al., 2010) gives us increasing confidence that these gregarine Apicomplexa are well-assembled with truly small genomes. Likewise, the consistently small set of protein-coding genes (∼2,200 – 2,600) raise questions about the core set of genes necessary for biological function in Apicomplexa. Reductions in genome size and gene content of parasitic species are well-known phenomena (Keeling, 2004; Keeling & Slamovits, 2005), but these three parasites of Lepidoptera apparently have half the gene content of even other arthropod-infecting gregarines in the genus *Porospora* (Boisard et al., 2022). More research is merited on this large-scale reduction in protein coding genes overall, but as a start, we have parsed basic patterns of gene conservation.

Across the three lineages of *Ophryocystis*, less than half of the gene content (993 genes) is strictly conserved. When accounting for the pairwise orthologous relationships, each lineage is left with <700 lineage-specific genes based on current analysis. Although these numbers are still too large to fully manually explore, additional data from related, yet un-sequenced parasites (Müller□Theissen et al., 2025) can only shrink these estimates. It may soon be tractable to explore the genetic basis of host-specificity of these parasites based on a relatively small number of candidate genes. For now, we explore already identified set of genes-of-interest.

### *ATPase evolution in* Ophryocystis

ATPases are class of proteins with diverse functions, but generally act to actively transport molecules either within or into and out of cells (Chène, 2002). For single-celled Apicomplexa important functions like creating ion gradients for cell-cell signaling (Kelly et al., 1985) are not relevant, but even single cells need to maintain proper salt balance and transport molecules against the concentration gradient. Some Apicomplexa possess ATP4 type sodium pumps (Lehane et al., 2019) but these have never been reported in *Ophryocystis* (Mongue et al., 2023); the *Helicoverpa*-infecting *Ophryocystis* apparently lacks this class of ATPases as well. Other gregarine Apicomplexa do possess calcium ATPases, both intracellular and membrane-bound (Lehane et al., 2019).

Mongue et al. previously reported a pair of PMCAs in milkweed butterfly infecting *Ophryocystis* albeit with especially long branches (2023). However, in our current re-analysis, we identify these ATPases as Type V phospholipid transporters, with a pair of orthologs in *Helicoverpa*- infecting *Ophryocystis*. This parasite has a more confidently assigned PMCA, closely related to PMCAs of other Apicomplexa and lacking in the two *Danaus Ophryocystis*. Although still not conclusive, the pattern produced by these new data and analyses is intriguing. Parasites of *Danaus* butterflies, which feed on plants rich in cardiac glycosides (Ackery & Vane-Wright, 1984), apparently lack any membrane-bound ion pumping ATPases. *Helicoverpa*, the host genus of this parasite, has a diet that does not include plants containing cardiac glycosides (Nasreen & Mustafa, 2000; Zalucki et al., 1986) and does retain a PMCA, like other gregarines (Mongue et al., 2023). Even though cardiac glycosides are not thought to pass through cell membranes, their inhibitory effects on cell membrane ion pumps has documented knock-on changes to intracellular calcium concentrations (Lee et al., 1970). Given that cardiac glycosides have some inhibitory properties for calcium pump ATPases as well (Kelly et al., 1985), it is possible that the loss of PMCAs in milkweed butterfly parasites is adaptive (Albalat & Cañestro, 2016), or at least neutral in a host environment where such ATPases would be routinely inhibited.

Finally, all three *Ophryocystis* have intracellular SERCA ATPases, though the branch length between *Helicoverpa* infecting *Ophryocystis* and the other two is longer than most others in this class of ATPases. The long branch length between SERCA ATPases of *Helicoverpa*-infecting *Ophryocystis* and the milkweed butterfly parasites may reflect an evolutionary history of changes to SERCA sequence in *Ophryocystis* prior to the loss of PMCA ATPases. This evolutionary history is still speculative and even if true does not explain the continued inhibitory effects of cardiac glycosides on *O. elektroscirrha* (de Roode et al., 2008). Nevertheless, the sequencing of another *Ophryocystis* genome has helped refine hypotheses on molecular mechanisms of host-parasite co-evolution.

## Conclusions

We have sequenced and annotated the genome of *Helicoverpa-*infecting *Ophryocystis*. Our results add a valuable resource to studying host-parasite dynamics. Comparisons with other sequenced *Ophryocystis* offer some additional clarity on adaptation to host biochemistry as well. More generally, these analyses begin to shed light on a poorly sampled portion of the apicomplexan tree of life and demonstrate the crucial role that genomics can play in resolving between organisms with few to no informative phenotypic differences.

## Data availability

Because this parasite species has yet to be formally described, it does not fit easily into frameworks like NCBI that require existing species metadata. Instead, raw sequencing reads, assembly, annotation, and orthology datasets are all available at https://doi.org/10.5281/zenodo.15856773.

